# Cas12a is a dynamic and precise RNA-guided nuclease without off-target activity on λ-DNA

**DOI:** 10.1101/2021.06.09.447528

**Authors:** Bijoya Paul, Loïc Chaubet, Emma Verver, Guillermo Montoya

## Abstract

Cas12a is an RNA-guided endonuclease that is emerging as a powerful genome-editing tool. Here we combined optical tweezers with fluorescence to monitor Cas12a binding onto λ-DNA, providing insight into its DNA binding and cleavage mechanisms. At low forces Cas12a binds DNA specifically with two off-target sites, while at higher forces numerous binding events appear driven by the mechanical distortion of the DNA and partial matches to the crRNA. Despite the multiple binding events, cleavage is only observed on the target site at low forces, when the DNA is flexible. Activity assays show that the preferential off-target sites are not cleaved, and the λ-DNA is severed at the target site. This precision is also observed in Cas12a variants where the specific dsDNA and the unspecific ssDNA cleavage are dissociated or nick the target DNA. We propose that Cas12a and its variants are precise endonucleases that efficiently scan the DNA for its target but only cleave the selected site in the λ-DNA.

## Introduction

<u>C</u>lustered <u>R</u>egularly Interspaced <u>S</u>hort <u>P</u>alindromic <u>R</u>epeats (CRISPR) that associate with <u>C</u>RISPR <u>as</u>sociated (Cas) proteins, constitute an adaptive immune system in bacteria and archaea against foreign mobile genetic elements (MGEs), such as plasmids and phages. These RNA-guided nucleases have been repurposed as versatile genome editing tools, triggering a technological revolution in the life sciences (Cong, 2013; Gasiunas, 2012; Jinek et al., 2012). CRISPR-Cas immunity maintains a genetic record of encounters with invading MGEs in the CRISPR array in the form of unique DNA segments (spacers), which are intercalated between short repetitive sequences (repeats). Transcription of the CRISPR array yields a long pre-CRISPR RNA (pre-crRNA), which after maturation into shorter crRNA molecules, forms a ribonucleoprotein (RNP) complex with the effector protein, resulting in a functional RNA-guided endonuclease responsible for interference. To maintain the integrity of the CRISPR array, the target DNA cleavage depends on the recognition of a short protospacer adjacent motif (PAM) sequence located upstream of the target site. The 3- to 4-nt PAM sequence recognized by the effector complex is present in the invading DNA molecules but not in the CRISPR array, therefore protecting this DNA region from cleavage (Mojica et al., 2009).

Depending on the type of RNP effector nucleases, CRISPR-Cas systems can be categorized into two classes (1 and 2), which are further subdivided into six types (types I through VI). Class 2 systems form an RNP complex between the multidomain effector Cas protein and a CRISPR RNA (crRNA) that contains the necessary information to target a specific nucleic acid sequence (Gasiunas, 2012; Horvath and Barrangou, 2010; Yan et al., 2018). Cas9 is the best characterized member of Class 2 (Type II-A) and it has been developed into a genome editing tool (Cong, 2013; Hsu et al., 2014; Jinek et al., 2012; Jinek et al., 2013; Kim et al., 2016; Kleinstiver et al., 2016; Zetsche et al., 2015). Diverse Cas9 nuclease variants are involved in clinical trials to develop new therapies against human diseases (Ernst et al., 2020). Recently, other members of Class 2 type V-A (Cas12a) and V-B (Cas12b, CasΦ, Cas12g, Cas14) have been studied (Harrington et al., 2018; Karvelis et al., 2020; Lee et al., 2019; Li et al., 2021; Ming et al., 2020; Pausch et al., 2020; Ren et al., 2021; Zhang et al., 2017b). However, they have not been as extensively characterized as Cas9.

Here we focus on Cas12a, this RNA-guided nuclease adopts a bilobed architecture with a recognition lobe (REC) and a nuclease lobe (NUC). In contrast with Cas9, which generates a blunt-DNA cleavage next to the PAM, Cas12a introduces a 6-7-nt staggered cleavage 70 Å away from the PAM sequence using a single catalytic site in a cleft between the RuvC and NuC domains (Yamano et al., 2016; Zetsche et al., 2015). The DNA overhang is produced by the different paths followed by the non-target (NT) and the target (T) strands to reach the catalytic pocket (Stella et al., 2017b). The absence of a HNH domain in Cas12a implies that the catalytic site in RuvC cleaves both the NT- and T-strands of the target DNA. However, while the NT-strand can reach the catalytic pocket with a 5′-3′ polarity, a conformational change in the distal side of the REC and Nuc lobes of Cas12a is needed to accommodate the T-strand with the proper polarity for cleavage. This results in the faster severing of the NT-compared to the T-strand (Stella et al., 2018a; Swarts and Jinek, 2019a). In addition to its specific endonuclease activity, an indiscriminate catalytic single-strand (ss) DNase activity was discovered in Cas12a. This secondary activity is triggered by binding of the target DNA or ssDNA complementary to the crRNA, thus mimicking the T-strand (Chen et al., 2018). Therefore, Cas12a is capable of precise cleavage of target DNA and indiscriminate degradation of ssDNA after activation. This activity has further expanded the possible applications of Cas12a to molecular diagnostics (Chen et al., 2018).

Despite its efficacy and versatility, the use of Cas9 in genome editing is limited due to toxic and possible mutagenic effects (off-target) (Cullot et al., 2019; Kosicki et al., 2018). These problems have boosted the search of CRISPR-Cas systems that can be developed into new genome editing tools. Cas12a is an attractive candidate for this purpose, specially from the standpoint of delivery via viral vectors due to its small crRNA (∼40nt) (Kim et al., 2017a; Kim et al., 2017b; Li et al., 2018; Zetsche et al., 2017; Zhang et al., 2017a), and its unique RNase activity, which enables Cas12a to process its pre-crRNA into mature crRNAs (Fonfara et al., 2016; Zetsche et al., 2015). This activity of Cas12a has been exploited for use in multiplex gene regulation and has been shown to successfully edit multiple endogenous targets simultaneously through delivery of multiple crRNAs on a single plasmid (Campa et al., 2019; Zetsche et al., 2017).

To understand Cas12a mechanics, we combined optical tweezers with confocal microscopy to dissect Cas12a binding and cleavage on force-stretched λ-DNA in real time. Our results show that Cas12a binds at different locations and its binding is more promiscuous at high forces; however, it only cleaves the selected target when low forces are applied to the λ-DNA, suggesting that it can work efficiently and precisely without substantial off-target events. We have also analyzed three different variants that present different cleavage activities and are potential candidates for biomedical applications. The mechanics of the optimized variants support the conformational model of Cas12a activation (Stella et al., 2018a).

## Results

### Cas12a displays dynamic binding and cleaves dsDNA

To monitor Cas12a-DNA interactions on the DNA at the single molecule level we combined optical tweezers with confocal fluorescence microscopy and microfluidics (Fig. 1A) (Gross et al., 2010; Heller, 2013). We designed 5 crRNAs targeting λ-DNA and tested the cleavage activity of the enzyme *in vitro*. The crRNA-1 displayed the higher activity, and thereby was selected for the study. Cas12a was assembled with a 3’-Cy5-labeled version of crRNA-1 to form the ribonucleoprotein complex. Binding along the λ-DNA was visualized by recording kymographs from the Cy5 fluorescent label at the 3’of the crRNA (Fig. 1B, Methods). We selected a crRNA sequence that binds a unique target site located at 29.7 kb on λ-DNA (Supplementary Table 1). The λ-DNA was attached to two polystyrene beads using the biotin-streptavidin affinity system and a force of 10 pN was applied to the λ-DNA pulling the beads. This force allows stretching of the λ-DNA without altering its base pairing (Smith et al., 1996b) (Zhang et al., 2013). In order to use the wild-type Cas12a and avoid cleavage of the target DNA, we performed binding experiments with the crRNA-1/Cas12a complex in the absence of Mg^2+^. The assay showed Cas12a binding on dsDNA without cleaving the tether (Fig. 1B). No binding was observed with 3’-Cy5-labeled crRNA-1 alone, confirming that the observed events represent binding of the assembled ribonucleoparticle (Fig. S1A). To visualize Cas12a cleavage, we included Mg^2+^ in the buffer. The presence of the divalent metal ion triggered an immediate loss of tension on the tether, indicating the disruption of the DNA molecule (Fig. 1C).

**Figure 1.**
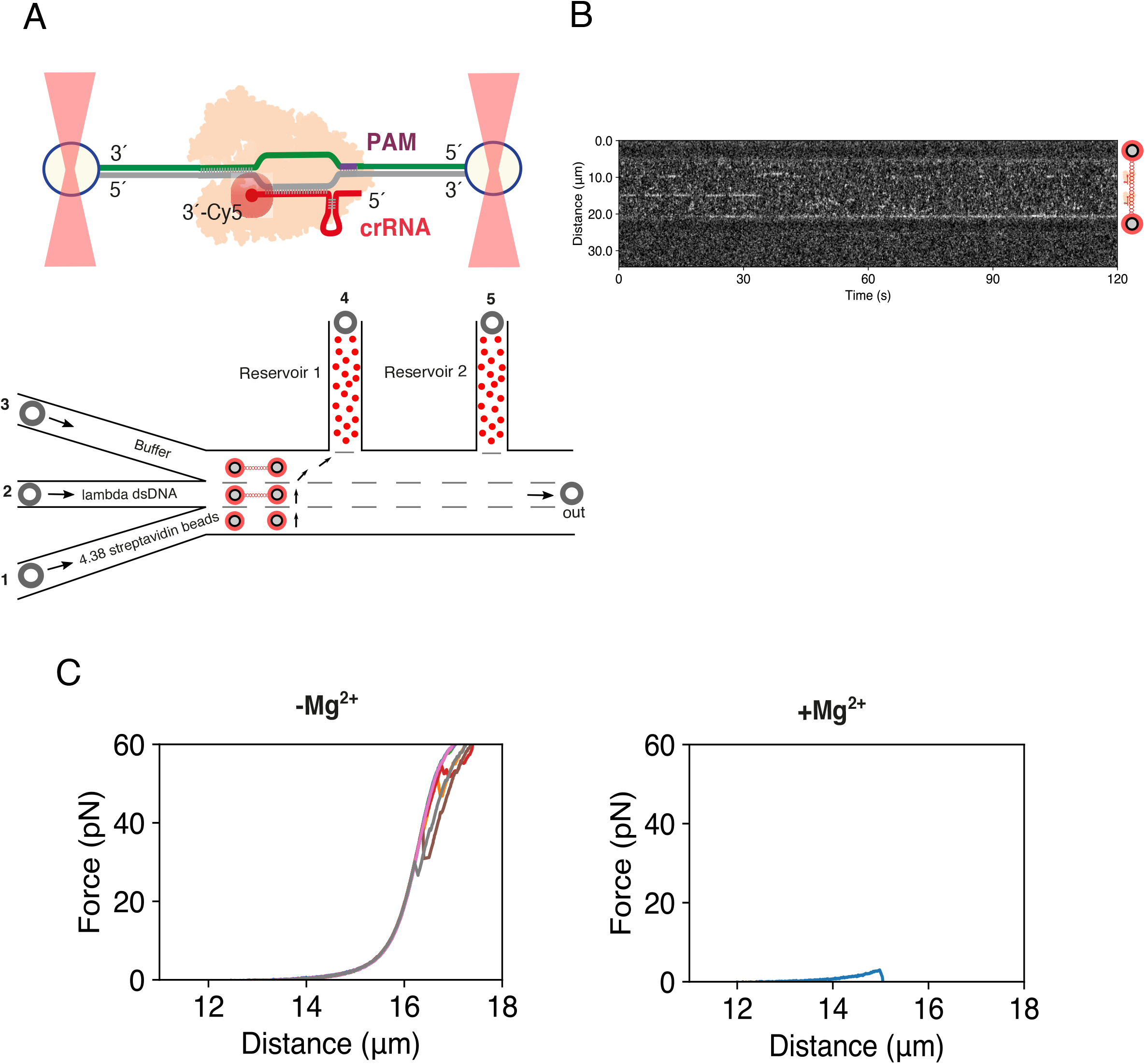
Experimental setup of Cas12a binding and cleavage onto λ-DNA. **A)** Schematic of assay (top) and schematic of microfluidic chip (bottom). **B)** Example of kymograph showing Cas12a binding (red channel). A single-pixel line scan is made along the DNA axis and plotted over time. **C)** Representative force-distance curves in the presence and absence of Mg^2+^. In the absence of Mg^2+^, the force increases with increasing inter-bead distance, while in the presence of Mg^2+^, the DNA tether ruptures at low forces, as marked by a sudden drop in force.

### Off-target Cas12a binding correlates with tension

Cas12a binding kymographs were also obtained at forces from 10 pN to 50 pN (Fig. 2A). The signal of the kymographs was averaged over a time-window to obtain fluorescent intensity profiles (Methods). The section of the kymograph corresponding to the DNA was identified using a bead detection algorithm and mapped to kbp (Fig. 2B, Fig. S1B-C, Methods). By using 10s-windows over the 120s-long kymograph, we obtained 12 intensity profiles per force, with each intensity profile showing the binding location and binding intensity averaged over a 10s window (Fig. 2C).

**Figure 2.**
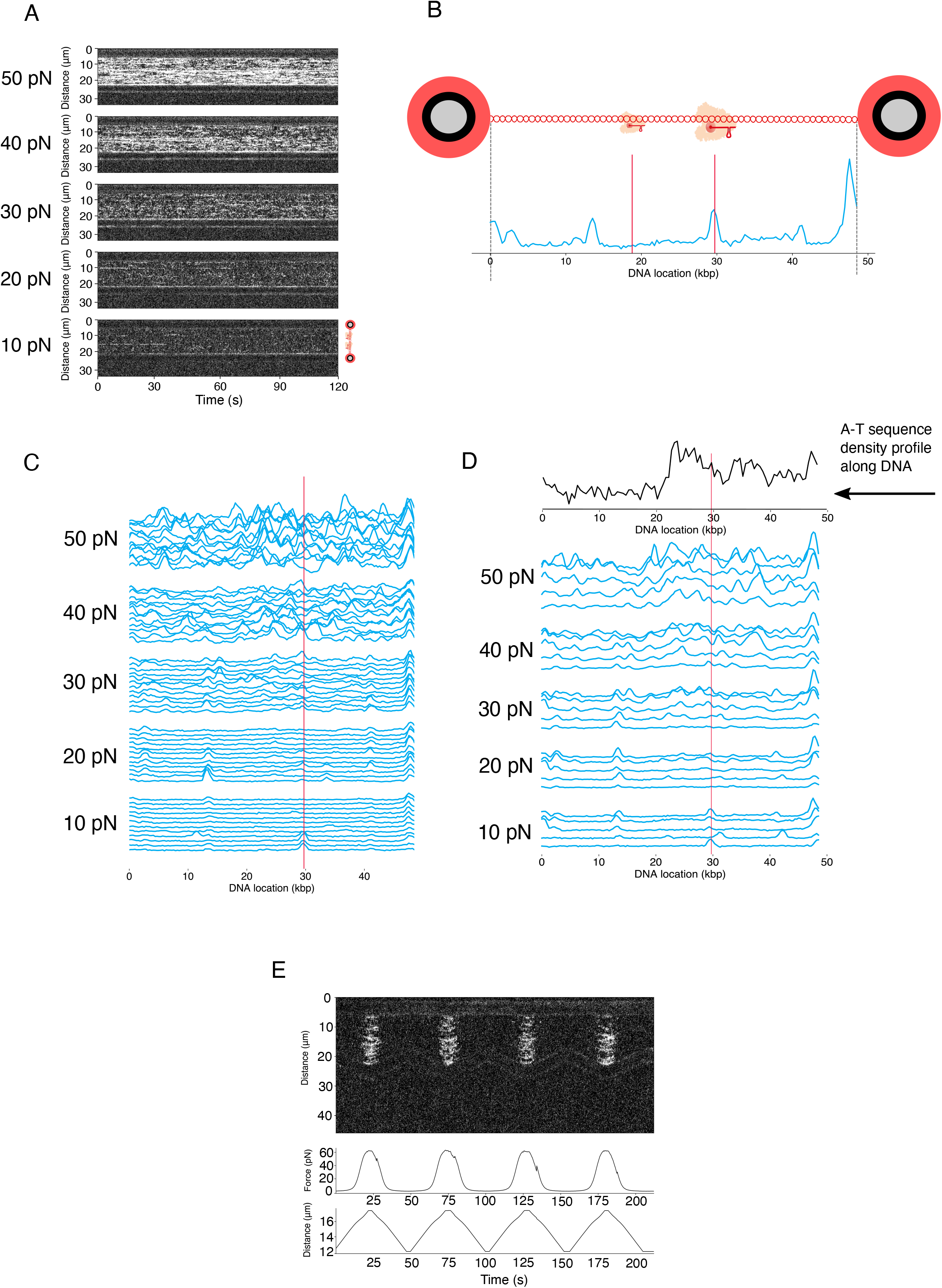
Kymographs and binding profiles of Cas12a with increasing tension. A) Representative series of 120s kymographs, showing Cas12a binding with increasing forces (n=5). **B)** Schematic showing how profiles were generated. Line-scans were time-binned and cumulative fluorescence intensity was plotted along the DNA axis. **C)** A single representative kymograph was binned 10s at each force, visualizing the dynamic nature of binding events. The red line indicates the target site centered at 29.707 kbps, which is at the middle of the target sequence including a TTTN PAM. **D)** Repeats of binding profiles at increasing forces binned at 120s (n=5). The red line indicates the target site centered at 29.707 kbps. The A-T density profile along the λ-DNA is indicated at the top of the plot. **E)** Representative kymograph and force trace of a repeated force-distance cycling (n=15).

In contrast with Cas9 (Newton et al., 2019), Cas12a showed a dynamic binding behavior and displayed both steady binding to λ-DNA over several 10s intervals and a more dynamic binding with shorter events when a 10 pN tension was applied on the DNA molecule (Fig. 1B, 2A-C). Sometimes after initial binding at a given site, the crRNA-1/Cas12a complex becomes no longer visible, suggesting that Cy5 could have been photobleached or that crRNA-1/Cas12a dissociates from the given site perhaps due to the absence of Mg^2+^, as it cannot complete cleavage. At high forces, between 30 to 50 pN, the increase of tension in the DNA molecule facilitates the binding of the ribonucleoprotein in multiple sites, indicating off-target binding events (Fig. 2C). Alternatively, by using a single 120s-long window, we obtained a single intensity profile per force (Fig. 2D). The resulting intensity profiles displayed different binding sites with an overall higher tendency to locate in the AT-rich regions of the λ-DNA (Fig. 2D), in contrast to Cas9 which shows a GC region preference (Newton et al., 2019). The correlation between increased tension and increased binding is highly reproducible. Indeed, reducing the force down to 10 pN results in the dissociation of most complexes from the DNA (Fig. 2E). This behaviour is in agreement with the PAM preference of *Francisella novicida* Cas12a (TTTN) and the PAM scanning stage of the proposed model of PAM-dependent DNA recognition and ATP-independent unwinding (Stella et al., 2017b).

### Cas12a off-target binding aligns with predicted PAM locations

Next, we investigated the location of the off-target sites by analysing the λ-DNA sequence. We searched for partial matches with the crRNA that could induce off-target binding with the crRNA, taking into account the PAM preference of Cas12a (TTN/TTTN). We searched for Watson-Crick coupling using short nucleotide sequences complementary to the crRNA (4-7 nt) starting at the seed sequence of the crRNA-1 spacer across the full λ-DNA sequence (Methods, Fig. S2).

If we do not consider the PAM as a condition *sine qua non* for binding, the analysis shows that there is a total of 251 different sites on λ-DNA that match with at least 4-nt of the crRNA sequence (including the target site) (Fig. S2A). However, PAM recognition is an indispensable preliminary event for Cas12a target DNA cleavage (Zetsche et al., 2015). Therefore, we performed a similar analysis considering the TTN and TTTN PAM sequences. While including TTN and 4-nts immediately after the PAM, the number of matching sites is reduced to 29 (Fig. S2B), and further reduced to 8 matching sites when using TTTN (Fig. 3A). Interestingly, the predictions of the off-target sites based on the PAM location and the adjacent sequence complementarity correlates strongly with the mapping of the binding profile (Fig. 3A, Fig. S2A-B), indicating that the consistent off-target binding of Cas12a is not random but directed by Watson-Crick coupling. Our sequence analysis of the λ-DNA also shows a few preferred off-target binding sites, one particularly consistent at 13.539 kbps which includes the TTTN-PAM and another nearby site at 13.153 kbps matched using the TTN-PAM (Fig. 3A)

**Figure 3.**
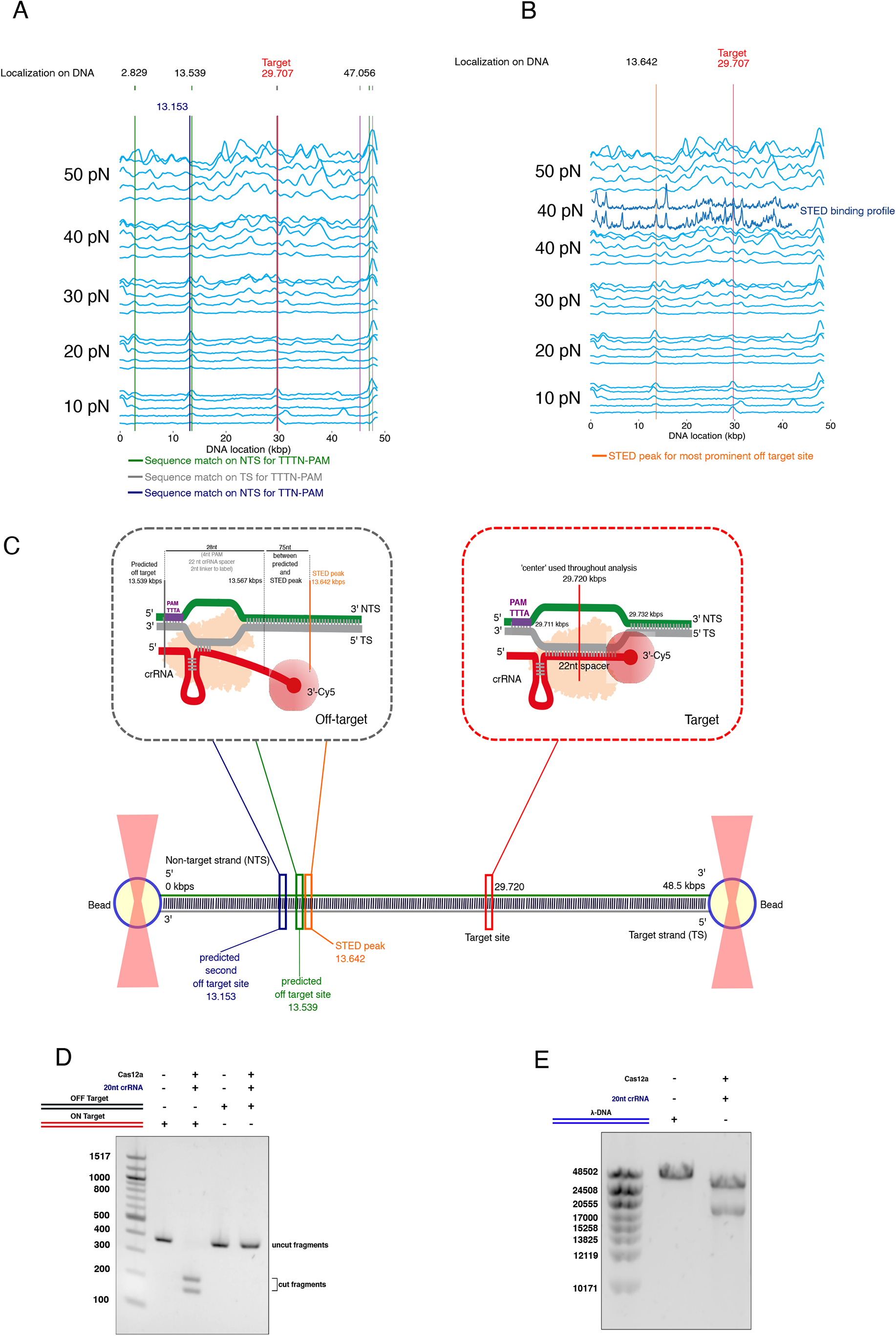
Analysis of predicted and observed off-target binding sites. **A)** Binding profile of Cas12a (120s binned, n=5) showing sequence matches to TTTN-PAM on NT-strand (5’→3’, green lines) and on T-strand (5’→3’, grey lines). The blue line indicates a single sequence match to TTN-PAM on the NT-strand (see Fig S2A-B for all matches). **B)** Binding profile of Cas12a (120s binned kymographs, n=5), including STED profiles at 40 pN (n=2). **C)** Scheme showing tentative positions of predicted and observed binding sites along the DNA. **D)** 2% agarose gel showing Cas12a enzymatic activity on 350-bp fragments of the λ-DNA containing the target and the off-target sites (n=3). **E)** 0.7% agarose gel showing the enzymatic activity of Cas12a on the complete λ-DNA (n=3)

We used Stimulated Emission Depletion (STED) microscopy at 40 pN to narrow down the two sequence matches at ∼13.5 kbp adjacent to the off-target binding peak, which were observed in the intensity profile of the confocal kymographs. This measurement was performed at 40 pN to enhance visualisation of off-target events and facilitate the localization of the main off-target site at 10-20 pN. STED showed individual binding events that were not resolvable by confocal imaging, and thereby not resolvable on the confocal intensity profiles (Fig. 3B). These binding events align well with the confocal intensity profiles. In addition, the STED intensity profile provides a better-defined experimental centre of the ∼13.5 kbp off-target site. After fine alignment of the STED intensity profile with the on-target peak (aligned at 29.707kbp, see Methods), the location of the preferential off-target binding site was identified at 13.642 kbp (Fig. 3C). Taking into account that there is at least 28-nt between the PAM and the Cy5 label on the 3’end (2-nt spacer, 22-nt crRNA guide, 4-nt PAM sequence, see Supplementary Table 1, Methods), there is only 75-nt difference between the experimental STED localization and the theoretical TTTN-PAM localization in the λ-DNA, thereby indicating a strong correlation between the two localisation techniques. The second predicted sequence match (with TTN PAM) found at 13.153 kbp shows less sequence homology and is 489-bp away from the STED-localized peak. Therefore, the resolution of the STED data suggests that Cas12a binds to the TTTN match at 13.539 kbp but not the TTN match at 13.153 kbp (Fig. 3C).

To evaluate whether this off-target binding event could lead to a DSB we performed a cleavage assay using a target DNA of 350-nt including the 13.539 kbp off-target binding site and tested the activity of the crRNA-1/Cas12a complex in the presence of Mg^2+^. In parallel, the experiment was performed with another 350-nt DNA containing the target site at 29.720 kbp (Fig. 3D, Supplementary Table 1). Cleavage could not be detected in the off-target binding sites, while the crRNA-1/Cas12a complex efficiently generated a DSB in the selected target DNA. To further evaluate whether any of the other binding events could lead to unspecific cleavage, we performed a similar activity assay but using the complete λ-DNA as substrate (Fig. 3E). The linear molecule was incubated with the crRNA-1/Cas12a in the presence of Mg^2+^ under the previous conditions. Cleavage was only observed at the selected site, indicating that despite the observed off-target binding activity, Cas12a only generates a DSB at the specific site.

Collectively, these results suggest that Cas12a performs an initial scanning of the possible PAM sites and tests complementarity with the guide (Stella et al., 2017b). However, the target DNA is severed only when the PAM and the subsequent crRNA/T-strand hybrid requirements have been fulfilled (Stella et al., 2018a). In contrast to Cas9 (Newton et al., 2019), Cas12a exhibits a dynamic binding at low forces, but cleavage is only observed at the target site.

### Cas12a cleaves λ-DNA at low forces

Next, we analyzed the specific cleavage of Cas12a and its differences with Cas9, we performed an optical tweezers experiment stretching the λ-DNA in the presence of Mg^2+^, thus allowing specific hydrolysis of the λ-DNA phosphodiester bonds (Stella et al., 2017b; Swarts et al., 2017; Yamano et al., 2016). The tension applied to the λ-DNA was progressively increased from 0 to 65 pN during 50s, after which the DNA molecule was relaxed back to lower force conditions. This cycle was repeated up to four times, and a tether breaking before the four cycles was considered a DSB initiated by the Cy5-crRNA-1/Cas12a complex. In several runs, where photobleaching had not occurred before cleavage, we were able to visualize the Cy5-crRNA-1/Cas12a complex bound to the cleaved DNA by stretching the cut DNA under flow (Fig. 4A). The break of the tether is detected by a sudden drop in force (Fig. 4B), and no DSB events were observed without Mg^2+^ (Fig. 1C). In contrast to Cas9, where a substantial mechanical force (40 pN) is needed to observe severing of the λ-DNA (Newton et al., 2019), DSB by Cas12a was exclusively observed below 5 pN of tension, either on the extension or relaxation steps (Fig. 4C), indicating that the DSB is not achieved when the DNA is stretched with high tension. Altogether, our binding and cleavage experiments suggest that while Cas12a efficiently explores possible target sites, phosphodiester hydrolysis to generate the DSB only occurs at the target site at low forces, suggesting that a high flexibility of the DNA is necessary to generate a DSB.

**Figure 4.**
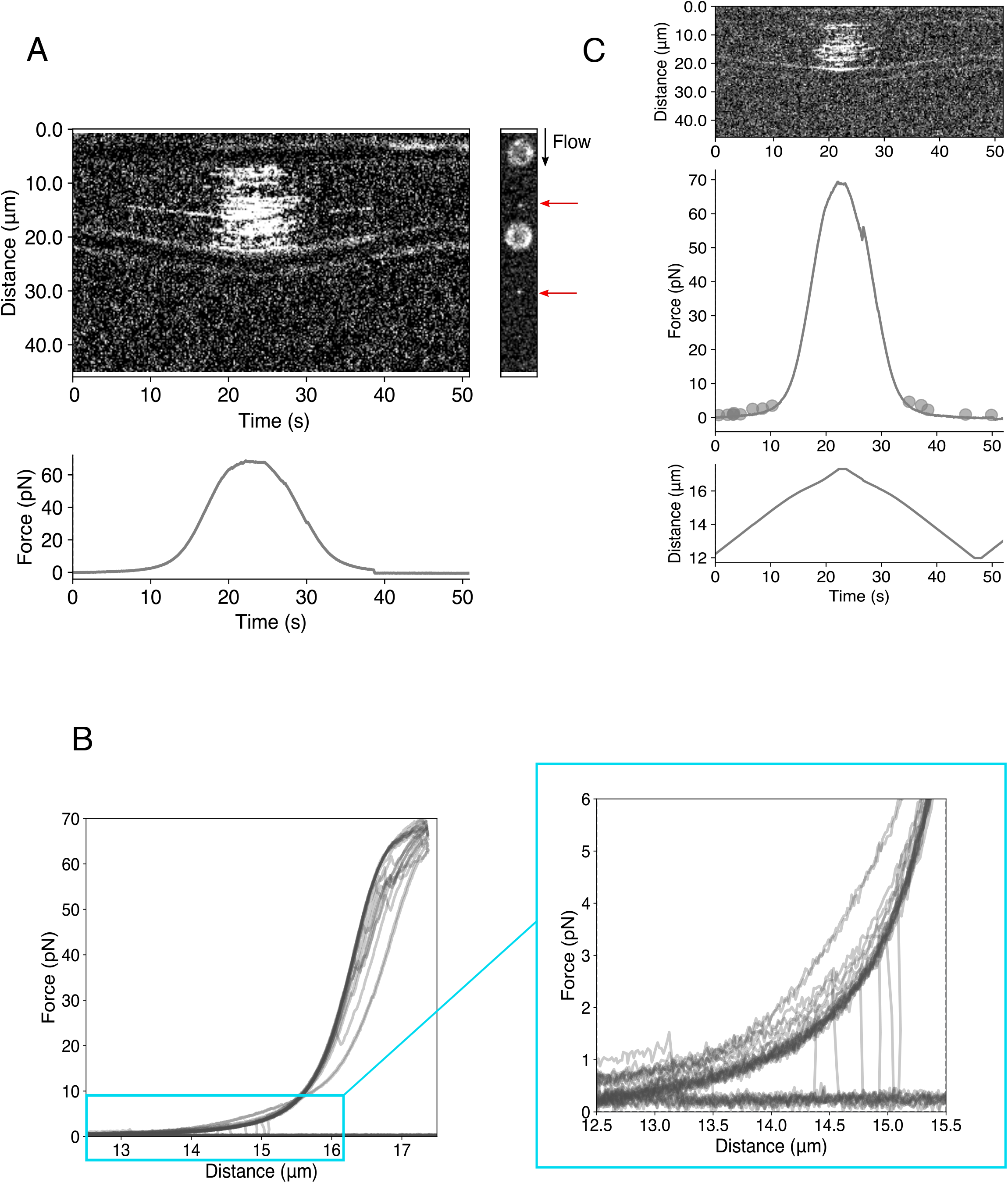
DSB occurs exclusively at low forces in the presence of Mg^2+^. **A)** Kymograph showing Cas12a binding during a force-distance cycle. The increment of inter-bead distance generates a increase in force applied to the tether. After relaxation, a DSB can be observed by the sudden drop in force. The right panel 2D scan shows Cas12a bound to the ends of the flow-extended DNA tether after DSB. **B)** Force-extensions curve of DNA tethers in the presence of Cas12a and Mg^2+^ (n=15). The inset depicts the sudden drop in tension indicative of a DSB generation. **C)** Summary of all DSB events measured in the presence of Cas12a and Mg^2+^ during force-distance cycles (n=14/15). The top panel shows a representative kymograph of Cas12a binding during force increase. In the middle panel, all DSB events (n=14, points) are plotted on a force-distance curve from the corresponding kymograph. All 14 DSB events happened at forces below 10 pN. 9 of them occurred on the force ramp up, while 5 were observed on the force ramp down. The bottom panel depicts the distance between the two beads *vs* time.

### Cas12a variants binding to dsDNA

After dissecting wild type (wt) Cas12a mechanical properties during target DNA recognition and cleavage, we analysed three engineered variants with different cutting properties. The fact that Cas12a displays an indiscriminate ssDNA degradation activity, which is observed in all its orthologs (Chen et al., 2018), together with recent studies that reported non-specific nicking of target sequences bearing mismatches in distal regions of the target DNA (Murugan et al., 2020), suggest that these cutting activities could be a problem for potential applications. To overcome these problems, we have engineered three variants of Cas12a with unique cleavage properties (Stella et al., 2018a).

The variants display different cleavage patterns in activity assays (Fig. 5A-B). Therefore, we dissected their binding and cleavage properties using the optical tweezers combined with confocal microscopy to understand the molecular differences with the wild type. These variants are: *(i)* the REC-Sub mutant (Cas12a_M1), which only displays unspecific ssDNA cleavage after activation with at least a 16-17 nt ssDNA complementary to the guide; *(ii)* the Q1025G/E1028G mutant (Cas12a_M2), whose ssDNA unspecific cleavage activity has been abolished and only cuts dsDNA specifically; and *(iii)* the K1013G/R1014G mutant (Cas12a_M3), which presents a specific nickase activity, as it displays preferential phosphodiester cleavage on the T-but not on the NT-strand of the target DNA. In addition, we included in the assays the E1006A-D917A catalytically dead mutant (Cas12a_KO) as a control (Fig. 5A-B).

**Figure 5.**
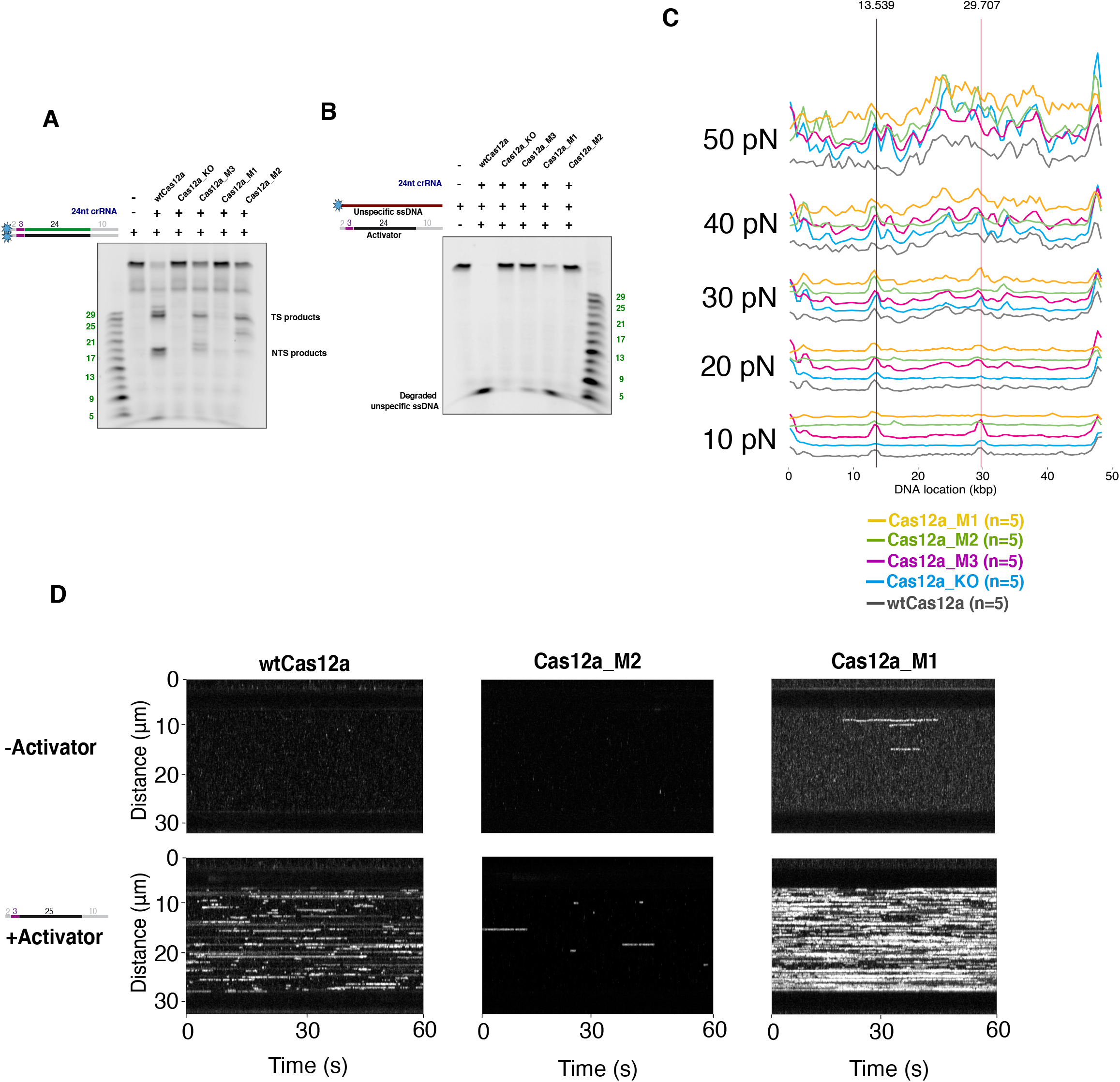
Cas12a variants biochemical cleavage assay and binding on dsDNA and ssDNA. **A)** 15% urea gel showing bulk dsDNA cleavage properties of wtCas12a and its variants (n=3). **B)** 15% urea gel showing bulk cleavage of indiscriminate ssDNA by wtCas12a and its variants (n=3). **C)** Binding profiles of wtCas12a and its variants at increasing forces binned at 120s (n=5, at each force for all the samples). The red line indicates the target site centered at 29.707 kbp, which is located at the middle of the target sequence including a TTTN PAM. The black line indicates the prominent off-target binding site at 13.539 kbp. **D)** Representative 60s-long kymographs showing binding of wtCas12a and its variants to ssDNA in the presence and absence of the activator at a force of 20 pN. All kymographs were obtained in the presence of Mg^2+^. In addition, in experiment the ssDNA tether was first exposed to a buffer without the activator, then, the same tether was moved to the buffer with the activator. In the case of wtCas12a and Cas12a-M1, the kymographs show a section of the data prior to the breaking of the tether.

The experiments were performed in the same conditions as in (Fig.2A, Methods). Overall, all the mutants show a binding behaviour to λ-DNA similar to wtCas12a, displaying only small variations between repetitive kymographs. The binding profiles depict an on-target site at 29.707 kbp and the main off-target site at 13.539 kbp at 10 pN (Fig. 5C, S3A). Besides these main sites, all of them also display a preference to bind the AT-rich region of the λ-DNA at high forces (Fig. S3A-B). Collectively, we observed that all the variants conserve a very dynamic binding to dsDNA which is comparable to the wild type, suggesting that their PAM scanning behaviour is not altered.

### Cas12a variants binding to ssDNA

As unspecific cleavage can be unleashed after specific DSB generation (Chen et al., 2018), we investigated in detail the binding of M1 and M2 on ssDNA, as we dissociated in these variants the specific dsDNA cleavage and the unspecific ssDNA degradation after activation (Fig. 5A-B) (Stella et al., 2018a). For that purpose, we tested binding to ssDNA using the λ-DNA strand corresponding to the “T-strand” as tether in the presence and absence of an activator of 40-nt at 20 pN (Fig. 5D, Supplementary Table I). For wtCas12a, we did not observe binding on ssDNA in the absence of activator. However, when the activator is present multiple binding events on the ssDNA can be recorded in the 60s kymograph, suggesting that assembly of the hybrid between the crRNA and the T-strand facilitates the interaction with the ssDNA. A similar behaviour can be observed for Cas12a_M1, which shows a slightly higher level of binding without the activator compared to the wild type but displayed a remarkably high number of binding events when the activator was present. This indicates that the mutations in Cas12a_M1 promote ssDNA binding in the presence of activator. On the other hand, the Cas12a_M2 mutant shows no binding to ssDNA without activator and a minimal binding when the activator is present. The observed Cas12a_M1 and Cas12a_M2 binding behaviour supports the bulk activity assays (Fig. 5A-B), showing that Cas12a_M1 preferentially cuts ssDNA while Cas12a_M2 is unable to unleash the unspecific ssDNA activity and therefore it also does not bind ssDNA.

Collectively the ssDNA binding behaviour of these variants supports the idea that the hybrid assembly between the crRNA and the activator promote the conformational change that opens the “lid” in the catalytic pocket (Stella et al., 2018a), allowing binding to the ssDNA to hydrolyse the phosphodiester bonds. By contrast binding on ssDNA is not observed for the Cas12a_M2, whose unspecific ssDNA cleavage activity has been eliminated by the double mutation and only cleaves dsDNA targets.

### Cas12a variants cleavage on ds- and ssDNA

To analyze the cleavage activity on ds- and ssDNA, we subjected the variants to two types of experiments: A dynamic assay in which the dsDNA tether was repeatedly stretched from 0 to 65pN for up to four times (Fig. 6A), and a static format in which the ds- or ssDNA tether was maintained at 20pN for 200s (Fig. 6B). The assays were performed in the presence or absence of activator depending on the experiment and Mg^2+^ was included in these experiments (Fig. 5A-B).

**Figure 6.**
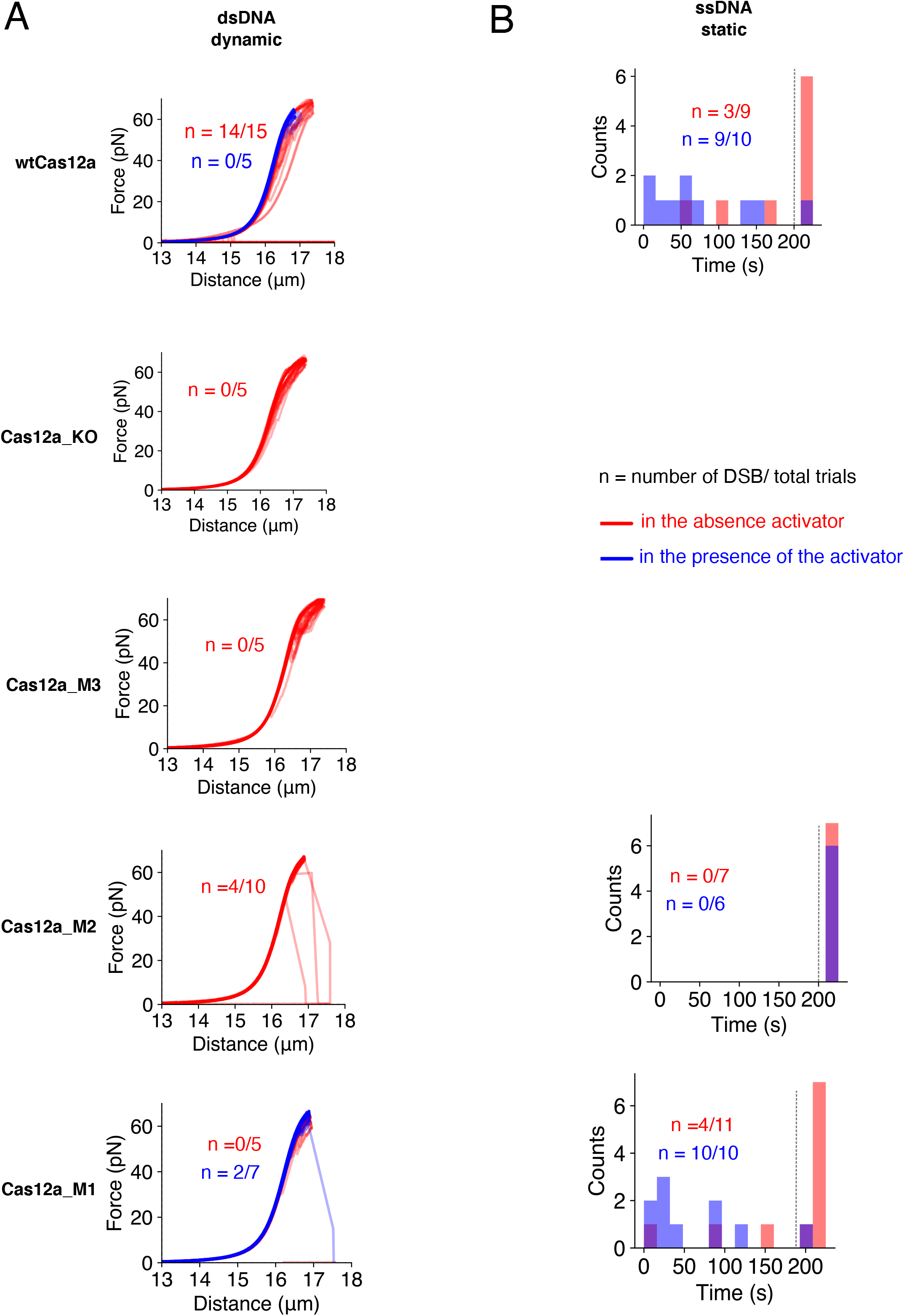
Quantitative analysis of dsDNA and ssDNA cleavage by wtCas12a and its variants. **A)** Force-extensions curves of λ-DNA tethers in the presence and Mg^2+^ and wtCas12a and its variants for dynamic dsDNA cleavage experiments. B) Plots of unspecific ssDNA degradation in a static setting by wtCas12a and its variants as a distribution of time. All experiments in the static form were conducted at 20 pN. λ-DNA ssDNA tethers that remained present after 4 force-distance cycles (A, dynamic) or 200s (B, static) were considered non-cleaved (indicated by the dashed line).

The generation of a DSB in the tether by wtCas12a is hampered by the presence of the activator. However, it enhances the cut on the ssDNA tether (Fig. 6A-B), suggesting that the formation of the hybrid between the crRNA and the activator disturbs the recognition of the target site on the λ-DNA and thereby cleavage. Yet the formation of the hybrid with the activator opens the catalytic RuvC/Nuc pocket, promoting the unspecific binding (Fig. 5D) and cutting of the ssDNA (Stella et al., 2018a).

Our experiments also confirmed that the Cas12a_M3 variant is unable to cleave dsDNA. No lack of tension was observed in the dsDNA tether when this mutant was tested (Fig. 6A-B). However, the use of Sytox orange, a DNA intercalator binding only to dsDNA (Biebricher et al., 2015; Yan et al., 2005), supported its “nickase” activity. The absence of this specific dye was used to detect a section of ssDNA, indicating a possible peeling of the DNA upon tension, and thereby suggesting that severing of one of the strands generated a region of ssDNA (Fig. S3C).

The binding analysis in the variants Cas12a_M2 and Cas12a_M1, where the unspecific ssDNA and specific dsDNA severing activities have been dissociated, show substantial differences. The bulk activities observed for these variants is well explained by our single molecule experiments. The dsDNA cleavage of Cas12a_M1 in the presence or absence of the activator is severely affected both in the static or dynamic experimental setup (Fig. 6A-B), as it was expected from its bulk activity and the binding experiments (Fig. 5B, 5D). By contrast to the wild type, Cas12a_M1 is able to partially cleave ssDNA in absence of the activator, while the presence of the complementary oligonucleotide triggers full activity, and all ssDNA molecules are severed within a wide distribution of time (Fig. 6B).

On the other hand, the ssDNA cleavage function is completely abolished in the Cas12a_M2, even when the activator is present. This variant solely cuts dsDNA, but its cleavage activity is reduced (Fig. 5A-B, 6A-B), suggesting that abrogation of the unspecific ssDNA activity affects the catalytic site. In contrast with wtCas12a, the DSBs produced by Cas12a_M2 on the λ-DNA were observed at high forces around 50 pN (Fig. 6A). The activity assay of Cas12a_M2 suggested that the T-strand is preferentially cut instead of the NT-strand (Fig. 5A). Therefore, to test this hypothesis, we performed an assay to determine whether the substitutions, besides eliminating unspecific ssDNA activity, reverted the cleavage preference of this mutant (Fig. S4). The assay showed that the Cas12a_M2 variant cuts the T-strand faster, suggesting that the redesign favours a conformation that disturbs phosphodiester hydrolysis in the NT-strand. This change in cleavage preference suggest that the tension in the DNA is necessary to permit the conformational change allowing NT-strand cleavage, and thereby visualisation of the dsDNA disruption. The preference to cleave the T-strand and not the NT-strand also suggests why this variant does not support unspecific ssDNA activity, we speculate that the entrance of the oligonucleotide would not be favoured by the conformation induced by the mutations.

Therefore, our single molecule experiments provide a mechanical background supporting the redesign in wtCas12a to dissociate and modify its ss- and dsDNA severing activities.

## Discussion

Currently, Cas9 is being tested in clinical trials to correct mutations or inactivate genes in different diseases using *ex vivo* approaches (Ernst et al., 2020). Despite its success, genome editing using RNA-guided nucleases is far from perfect and presents limitations. One of the main restrictions stems from the precision and ability of the nuclease to target and generate the break at the desired site (Cullot et al., 2019; Fu et al., 2013; Hsu et al., 2013; Kosicki et al., 2018; Lin et al., 2014). Structure-function studies of Cas12a have provided important snapshots of the molecular details of Cas12a (Stella et al., 2017b; Stella et al., 2018a; Swarts and Jinek, 2019b; Swarts et al., 2017; Yamano et al., 2017). Three regions, termed the “linker”, the “lid” and the “finger”, build a network of checkpoints along the main cavity of the protein that monitor the assembly of the hybrid between the crRNA and the T-strand to open the catalytic site in the RuvC/Nuc pocket of Cas12a (Stella et al., 2018a). Yet a complete characterization at the single molecule level of the target recognition and cleavage by Cas12a and its engineered variants to fully understand this novel endonuclease was lacking.

### Cas12a vs Cas9

Our study reveals that Cas12a displays a very dynamic binding behaviour on the λ-DNA (Fig. 2A). The ribonucleoprotein actively searches for the target sites by binding to different locations where a complete or cryptic PAM sequence (including partial matches with the crRNA) is found. The number of off-target sites observed for Cas12a increases when tension is applied to the λ-DNA (Fig. 2A-C). Two main binding sites are observed in the DNA (Fig. 1B, 2A). Overall, the lifetime of the binding events is short, but these events are longer and more persistent in the target site and off-target site sharing the TTTN PAM and 4-nt of the guide with the target site (Fig. 3A). After identifying the preferential off-target site, the cleavage experiments showed that binding of Cas12a did not lead to cleavage of a 350-nt dsDNA containing the cryptic off-target site. Furthermore, when a similar activity assay was performed with the complete λ-DNA a single cut corresponding to the selected target site could be observed. To understand this behavior, we analyzed Cas12a cleavage on the λ-DNA in the presence of Mg^2+^. The experiments revealed that Cas12a generates a DSB in the DNA at low forces between 2-4 pN (Fig. 4A, C), suggesting that Cas12a is unable to hold the ruptured DNA together.

The binding and cleavage behavior of Cas12a on λ-DNA are rather different than that of Cas9 (Newton et al., 2019). Like Cas12a, Cas9 off-target is enhanced when tension is applied to the DNA. However, in the case of Cas9, the force increase on the λ-DNA has been shown to induce off-target binding that promotes cleavage at 40 pN in multiple sites(Newton et al., 2019). By contrast Cas12a induces the break on the λ-DNA catalyzing the phosphodiester hydrolysis at low forces below 5pN (Fig. 4A, C). Although Cas9 and Cas12a are related molecules and share some degree of structural organization (Stella et al., 2017a), the two RNP complexes are quite different. In Cas9 cleavage occurs next to the PAM and there are two catalytic sites, one in the RuvC and other in the HNH domain. However, in Cas12a the PAM is 70 Å away from the single RuvC/Nuc pocket where catalysis occurs. This structural organization suggest that in Cas9, PAM recognition and catalysis of the blunt end DSB are closely associated, while in Cas12a PAM recognition is physically apart from the single RuvC/Nuc active site to generate the overhang. Hence, the event of PAM scanning and testing of complementarity is separated from the active site and needs the completion of at least a 16-17 nt long crRNA/T-strand hybrid to activate catalysis (Stella et al., 2018a). Furthermore, as RuvC domains preferentially cut DNA with the 5’-3’polarity (Wyatt and West, 2014), we have proposed that a conformational change should occur after the cleavage of the NT-strand to position the T-strand, which is in the 3’-5’ polarity coupled with the crRNA, in the RuvC catalytic pocket for cleavage (Stella et al., 2018a).

The dynamic binding of Cas12a revealed by our data supports the proposed PAM-dependent DNA recognition and unzipping model (Stella et al., 2017b), where Cas12a would bind PAM sequences and test complementarity with the crRNA before cutting the target DNA. In addition, the fact that Cas12a only generates the DSB in the λ-DNA at low forces, supports the notion that a conformational change is needed to position the T-strand in the catalytic pocket (Stella et al., 2018a). This conformational change would be hindered at high forces, as on a high-tension DNA Cas12a would be unable to “push” the stretched T-strand for phosphodiester hydrolysis into the RuvC/Nuc pocket with the proper polarity. Therefore, a DSB generated by Cas12a should be visualized at low forces (Fig. 4A, C). This is not the case for Cas9, as each strand is severed by a different catalytic center (Jinek et al., 2012). Therefore, the numerous off-target binding events of Cas9 at high forces combined with the stretched DNA conformation would favor its off-target behavior on λ-DNA (Newton et al., 2019). In summary, although Cas12a is able to bind numerous cryptic-sites, the conformational restrictions due to its single catalytic pocket avoid that these off-target binding events observed at medium and high forces (Fig. 2A-C) result in unspecific breaks on the DNA (Fig. 4A-C).

### Cas12a variants

The analysis of the three Cas12a variants shows that they conserve the dynamic binding behaviour to dsDNA (Fig. 5C, Fig. S3A-B). The off-target binding of the variants increased when force was applied to the λ-DNA and all of them show a more persistent binding to the target and display the same preference as the wild type for the off-target sites. This short-lived binding and the longer lifetime on the target and cryptic target sites indicate that the PAM scanning and initial unzipping mechanism to test hybridisation are not affected by the changes introduced in these mutants. However, the variants display a different cleaving capacity. The mutations are located in the active site in the Cas12a_M2 and Cas12a_M3 mutants, while for Cas12a_M1 they reside in the “linker”. Both the wild type and the Cas12a_M2 cleave dsDNA, while the Cas12a_M1 shows a minor DSB generation at high forces that may be due to its action on the ssDNA generated by the tension. We have eliminated the indiscriminate ssDNA cleavage in the Cas12a_M2 variant by inverting wtCas12a cleavage preference (Fig. S4). By doing so, the double mutant seems to induce a conformation that avoids severing ssDNA non-specifically. Therefore, this variant could potentially eliminate possible sources of off-targets in genome editing applications. However, the deletion of this collateral activity of Cas12a decreases the efficiency of the specific cleavage activity on dsDNA and an optimization of the catalysis is needed to improve its efficiency.

The Cas12a_M3 binds to dsDNA but it does not generate a DSB on the target and it does not display indiscriminate ssDNA activity (Fig. 5A-B). However, this variant is a sluggish enzyme as cleavage is not observed throughout the 200s of the dynamic experimental setup. This observation also supports the model suggesting that for severing the T-strand a conformational change is needed to locate the DNA in the active site. The activator-mediated cleavage of ssDNA in Cas12a and Cas12a_M1 correlates well with the observed binding and cleaving activity. Binding to ssDNA and cleavage is enhanced, especially in Cas12a_M1, in the presence of the activator in agreement with the activation model of catalysis after hybrid assembly (Stella et al., 2018a). In addition, cleavage shows a wide distribution of time, supporting our cleavage model and the unspecific character of this activity.

Our work shows that the combination of mechanical information with structural insights can render valuable information to redesign CRISPR-Cas endonucleases. In addition, this study confirms the molecular basis of the conformational activation of Cas12a catalysis after target DNA recognition. This study provides a biophysical background to understand the activity and precision of Cas12a and its variants in cellular and *in vivo* genome editing approaches

## Acknowledgements

The Novo Nordisk Foundation Center for Protein Research is supported financially by the Novo Nordisk Foundation (Grant NNF14CC0001) and the Distinguished Investigator (NNF18OC0055061) grants to G.M. Guillermo Montoya is member of the Integrative Structural Biology Cluster (ISBUC) at the University of Copenhagen. We thank Janna Bogers (LUMICKS) for her helpful assistance in C-Trap experiments. We thank N. Hatzakis for careful reading of our manuscript.

## Author Contributions

B.P. prepared the samples and performed biochemical and single molecule experiments together with L.C and E.V. The data were analysed by E.V., L.C. and B.P. The global results were discussed and evaluated with all authors. G.M. develop the concept, supervised the project and wrote the manuscript with input from all the authors.

## Declarations of interest

Guillermo Montoya declares that is a co-founder of Twelve Bio

## Supplementary Figures

**Figure S1. Controls for single molecule experiments, mapping of binding profiles to A-T density along λ-DNA and mapping of binding profile to base pairs along λ-DNA.** A) 3 kymographs of a single DNA tether which was exposed to 2nM crRNA-1-Cy5/Cas12a (top), then 2nM crRNA-1-Cy5 crRNA only (middle) and finally back into 2nM crRNA-1-Cy5/Cas12a (bottom), all at 50 pN. The binding observed in the top and bottom kymographs is specific to Cas12, as the labelled crRNA alone does not yield any binding. B) Representative kymograph illustrating the bead center, which is marked by a dot in the blue channel. This reference was used for localization. C) An intensity profile was generated along the entire length of the scanned λ-DNA.

**Figure S2. Sequence analysis for predicted binding locations**. **A)** Binding profile of Cas12a showing sequence matches without any PAM considerations on NT-strand (5’→3’, green lines) and on T-strand (5’→3’, grey lines). **B)** Binding profile of Cas12a showing sequence matches TTN-PAM on NT-strand (5’→3’, green lines) and on T-strand (5’→3’, grey lines). **C)** Scheme showing example search for a sequence match along λ-DNA.

**Figure S3. Binding of Cas12a variants to** λ**-DNA. A)** The top panel shows representative 120s kymographs of Cas12a variants with increasing forces (n=5). The bottom panel shows the binding profiles of Cas12a variants at increasing forces binned at 120s (n=5). **B)** Binding profiles of wtCas12a and its variants binned at 120s plotted alongside A-T density profile of the λ-DNA. **C)** Kymograph showing the increase of Cas12a-M3 binding events with force. The λ-DNA was co-stained with Sytox orange as a dsDNA marker.

**Figure S4. Kinetics of Cas12a_M2 cleavage on T-strand and NT-strand**. **A)** 15% urea gel showing the time-course cleavage assay of the T-strand and NT-strand (n=3). **B)** Quantification and apparent rates of product formation of the T-strand (grey) and NT-strand (green). The experiment was repeated 3 times. The error bars show the SD of each measurement.

## Methods

### Plasmid preparation

All experiments in this manuscript were performed with the *Francisella novicida* (Fn) Cas12a gene. All cloning and site directed mutagenesis were performed as previously described by Stella, Mesa et al., 2018 (Stella et al., 2018b).

### Protein expression and purification

The target construct containing pET21-FnCas12a was transformed into *E.coli* BL21 star (DE3) cells carrying the pRare2 plasmid (that provides seven tRNAs recognizing rare codons). A single colony containing both plasmids was selected to inoculate a culture of 50ml LB media, with 50 μg/ml ampicillin and 25 μg/ml chloramphenicol, grown overnight at 37°C, which was then used to inoculate 1L fresh LB media containing the same quantities of antibiotics and grown at 37°C until an OD_600_ between 0.7-0.9 was reached. The large-scale culture was then induced with 0.5mM isopropyl β-D-1-thiogalactopyranoside (IPTG) and grown at 37°C for an additional 3h. The cells were then harvested, resuspended in lysis buffer [50mM HEPES pH 7.5, 2M NaCl, 5mM MgCl_2_, 0.5mM TCEP, 1 tablet of complete Inhibitor cocktail EDTA free (Roche), per 50ml, 5 U/ml Benzonase, 160μg/ml lysozyme], and lysed by sonication for 10 mins with 15s ON and 20s OFF cycle. The lysate was then centrifuged and filtered to remove cell debris and insoluble particles, and then loaded onto a 5ml crude HisTrap^TM^ FF column (Cytiva) equilibrated in 50mM HEPES pH 7.5, 500mM NaCl, 10mM Imidazole, 0.5mM TCEP. The protein was eluted through a step gradient of 3%, 8% and 50% of the following buffer [50mM HEPES pH 7.5, 500mM NaCl, 500mM Imidazole, 0.5mM TCEP]. Enriched protein fractions from the 50% elution were loaded onto a 5ml HiTrap Heparin HP column (Cytiva) equilibrated with 50mM HEPES pH 7.5, 100mM NaCl, 0.5mM TCEP. Elution was performed through a linear gradient of 0-100% of the following buffer [50mM HEPES pH 7.5, 1M NaCl, 0.5mM TCEP]. Enriched protein fractions from the elution were concentrated and loaded onto a HiLoad Superdex 200 16/60 column (Cytiva) equilibrated in 50mM Bicine pH 8.0, 150mM KCl, 0.5mM TCEP. Protein rich fractions were concentrated using 50kDa Amicon® Ultra-15 Centrifugal Filter Units, flash frozen in liquid nitrogen and stored at -80°C.

### crRNA design for single molecule experiments

The region 29.711 kbps to 29.732 kbps on the Enterobacteria phage λ DNA was chosen as the prospective target because it contained the FnCas12a PAM sequence (TTN). The target region (spacer) of 22nt was placed on the 3’ end of the repeat sequence 5’ - AAUUUCUACUGUUGUAGAU-3’ to form a 41nt long crRNA-1. The crRNA was purchased from IDT, with fluorescent labels on the 3’ end (Cy5), 3’-Cy5-labeled crRNA-1.

### Bulk cleavage assays

The purified protein was incubated with the crRNA-2 (rArArU rUrUrC rUrArCr UrGrUr UrGrUr ArGrAr UrGrAr GrArA rGrUr CrArU rUrUrA rArUrA rArGrGr CrCrArCrU) in the molar ratio of protein:RNA 1:1.3, at room temperature for 15mins to form the ribonucleoprotein complex (RNP) in reconstitution buffer A with the following composition: 20mM Bicine pH 8.0, 150mM KCl, 5mM MgCl_2_. The fluorescently labelled dsDNA target was prepared by annealing the labelled target (CGA TGC ATG CAG TGG CCT TAT TAA ATG ACT TCT CTA ACG/36-FAM/) and non-target strands (/56-FAM/CGT TAG AGAAGT CAT TTA ATA AGG CCA CTG CAT GCA TCG) according to the manufacturer’s instructions (both oligos were independently purchased from IDT). The RNP from the former reaction was then incubated with the DNA substrate in a molar ratio of RNP:DNA 1:1.7, at 37°C for 30 minutes, and then were stopped after incubation by adding equal volumes of a stop solution (8M Urea and 100mM EDTA pH 8.0), followed by incubation at 95°C for 10 minutes. The stopped reactions were then loaded onto 15% Novex TBE-Urea Gels (Invitrogen) and run according to the manufacturer’s instructions and imaged using an Odyssey FC Imaging System (Li-Cor).

### Ratio of cleavage of the target and non-target strands

The RNP was prepared as described as above in the bulk binding assays, and then incubated with the annealed dsDNA substrate, also described above, in the same molar ratio. The reaction was monitored at several time points at 1, 5, 10, 30, 60, 90, 120 and 180 minutes by taking the reaction at those time points and stopping it with equal volumes of stop solution as described above, followed by incubation at 95°C for 10 minutes. The samples (containing 4 pmoles of dsDNA substrate) were then loaded onto 15% Novex TBE-Urea Gels (Invitrogen) and run according to the manufacturer’s instructions and imaged using an Odyssey FC Imaging System (Li-Cor). The intensity (I) of the DNA bands was quantified using ImageStudio. Considering the (I)_t=0_= 4 pmoles, the formation of products at different time points was calculated using the following formula pmoles_t=n_= 4 pmoles * (I)_t=n_/(I)_t=0_, where n is the different time points. The average of 3 independent experiments was used in the plot.

### Complex formation for single-molecule experiments

For optical tweezers experiments, Cas12a protein was complexed 1:1 with 3’-Cy5-labeled crRNA-1 (rArArUr UrUrCr UrArCr UrGrUr UrGrUr ArGrAr UrUrGr UrArA rArGrA rArArC rArGrUr ArArG rArUrA rArU/3Cy5Sp/) in a final concentration of 5uM and low volume of 20ul in buffer A (20mM Bicine pH 8.0, 150mM KCl, 5mM MgCl_2_). After adding the Cas12a protein and crRNA-1 together, the sample was left at RT for 15min after which it was kept on ice until further processing. For the experiments including the activator oligo, the Cas12a-crRNA complex was incubated with λ-activator, a 40 nt activator ssDNA (CAT CGG GTT GAG TAT TAT CTT ACT GTTT CTT TAC ATA AAC) at a ratio 1:2, 10uM final concentration, for 15-30min at 25°C, after which the sample was kept on ice. The complex stock solution was stored overnight at 4°C and subsequently used. For optical tweezers experiments, the 5uM complex stock solution was diluted in either buffer A (20mM Bicine pH 8.0, 150mM KCl, 5mM MgCl_2_) or buffer B (20mM Bicine pH 8.0, 150mM KCl), depending on whether the complex would be used for cleavage or binding measurements respectively, to a working concentration of 1nM (1:5000).

### Optical tweezers assay

Single-molecule experiments were performed at room temperature using a correlative confocal fluorescence - optical tweezers microscope (LUMICKS, C-Trap). Two traps were generated by a 1064nm laser passing through a 60x Water Immersion (NA1.2) objective. One trap was beam-steered by controlling a piezo mirror, while the other trap provided force measurements. Force measurements were performed by back-focal plane interferometry using a position-sensitive detector. The combination of micro-stage movements with a 5-channel microfluidics system allowed rapid in situ construction and examination of dumbbell assays.

### Surface passivation

To ensure a stable concentration, the channels harbouring Cas12-crRNA complex (“protein channel” hereafter) were passivated in multiple steps. After completion of the cleaning protocol, the protein channels were incubated with 0.1% BSA (in MQ) for 20min under flow. Then, the channels were incubated for 20min in 0.5% Pluronics (in MQ), again under flow. Finally, at least 250ul of the [1nM] complex working solution was loaded in the protein channels and left in the system ON. The next day, the complex working solution was replaced with a fresh working solution before starting experiments.

### Dilution and Oxygen Scavengers

On the day of experiments, a fresh working solution was made from the complex stock solution that was prepared the day before by diluting either in Buffer A (for cleavage experiments) or in Buffer B (for binding experiments and wildtype cleavage control with no Mg^2+^ ions) to a working solution of 1nM (1:5000). This working solution was supplemented with fresh Oxygen Scavengers (0.65% Glucose (D-(+)-Glucose, Sigma) (Pelz et al., 2016); 165U/ml Glucose Oxidase (Glucose Oxidase from *Aspergillus niger*, Sigma)^[2]^; 2170U/ml Catalase (Catalase from bovine liver, Sigma)(Roy et al., 2008)). The final working solution was replaced with fresh buffer every 90min.

### DNA templates & Beads

As a template, λ-DNA (48.5kb) was used containing biotin labels (LUMICKS, Amsterdam). For experiments using dsDNA templates, λ-DNA was labelled with biotin labels on complementary strands. For experiments using ssDNA, λ-DNA contained biotin labels on either side of the same strand, allowing the other strand to be melted off at higher tension (Candelli et al., 2014). DNA molecules were captured between two trapped streptavidin-coated polystyrene beads ( ?4.38 µm, Spherotech).

### Data acquisition

#### Workflow Binding assays

Beads were optically trapped and calibrated using the power spectrum method (Berg-Sørensen and Flyvbjerg, 2004). Next, DNA was tethered between the two beads by use of the multichannel laminar flow cell. The presence of a single DNA molecule was confirmed by comparing the force-distance curves to the inherent mechanical force-extension expected for single dsDNA (Smith et al., 1996a). In the case of ssDNA experiments, dsDNA molecules were stretched to high forces, causing the DNA to melt (Candelli et al., 2014).. For binding assays, the construct was subsequently moved to the protein channel. The applied force on the DNA molecule was increased in a stepwise manner by increasing the distance between the beads, maintaining at least 120sec at each force level. For all binding assays (all mutants), the experiments were performed in the absence of Magnesium.

#### Workflow Cleaving assays

Constructs were generated as described above. All experiments were performed in the presence of Mg^2+^ ions, except for the wild type control with no Mg^2+^. For static cleavage experiments, constructs were moved into the protein channel under a constant force of 20pN and kept there for at least 200sec. For dynamic cleavage experiments, constructs were moved into the protein channel at low force, after which a force-extension curve was initiated and repeated for at least 4 times (taking ±200s) or until cleavage occurred. Cleavage is defined as a loss of tether, marked by a sudden decrease in force. (Fig. 1C, 4B).

#### Fluorescence imaging

Fluorescence microscopy was achieved by imaging the construct with a confocal system using 532nm (for bead visualization) and 638nm excitation (for Cy5) lasers and corresponding Avalanche Photodiodes (APDs). Confocal pixel size was set to 100nm and the pixel dwell time was 0.05ms. For kymographs, a single pixel height line scan along the DNA axis was scanned and plotted over time (y=DNA molecule, x=time).

### Data analysis

#### 120s binding profile (binding location)

To analyse binding profiles (“profiles” hereafter) of Cas12a complexes on DNA, we obtained the mean fluorescence signal over a 120s time window (Fig. 1B). To map the binding of Cas12 to base pair, the naturally occurring signal in the 488 channel (Fig. S1C) from the polystyrene 4.38 μm beads that indicate the bead’s center was localized using a Gaussian filter. The center position was tracked across the whole 120s window. The signal coming from the DNA tether region (Fig. S1C) was defined as the bead center-to-center distance minus twice the radius of the beads. The DNA’s 48.5kb was then mapped to the DNA region. The background was defined as the minimal signal found in the region outside of the construct (Fig. S1C). Once the background was subtracted and the DNA distance was mapped in basepairs, a manual inspection of the binding profile was done to assess DNA orientation. The orientation of the DNA was determined based on alignment of the profile with the on-target site, recurrent off-target sites and with the typical asymmetrical binding preference that was consistently observed at higher forces. To facilitate data visualization and comparison between different kymographs, all profiles from a single protein were plotted on the same plot (grouped by force level). Each binding profile was further normalized to the highest intensity value present within any of the profiles. To pool all conditions into a single figure, an average was calculated over 500bp windows at each force for each protein. These profiles were then plotted against the other forces and conditions in a single plot. To facilitate data visualization, each of these profiles were further normalized to the highest intensity value present within these profiles. For more information on how the DNA-basepair mapping is done with an example data set, the analysis script is available on Harbor (www.harbor.lumicks.com).

#### 10s binding profile (binding dynamics)

In order to visualize binding dynamics, a single representative kymograph was selected. The 120sec kymograph was averaged in 12x 10sec windows to allow detection of short timescale dynamic events. Profiles were grouped per force and plotted consecutively to show dynamic changes over time. Each profile was normalized to the highest intensity value present within profiles across the full force range.

#### Cleavage

Cleavage events were captured using dynamic or static assays. In both assays, the sudden decrease in force within the 200s timeframe was manually identified through visual inspection. When no cleaving was observed before the 200s timeframe, these cases were classified as non-cleaving. Force-extension curves showing the sudden decrease in force were obtained by plotting the force in the x-direction on Trap 2, downsampled to 15 Hz (from 78 kHz) against the camera-based distance tracking. The time-to-cleavage distribution for the WT, Cas12a_M1 and Cas12a_M2 on ssDNA was obtained using an automated change-point analysis detecting the sudden drop in force.

#### Off-target data analysis

The theoretical off-target data analysis included searches for short nts sequences complementary to the crRNA, from 4 to 7 nts starting at the PAM-proximal side of the crRNA selected and compared across the full λ-DNA sequence. A match on the λ-DNA is found if anywhere on the DNA, on either strand, the short sequences are identical and in the same order. Short sequences including the TTN PAM (7-10 bps) and the TTTN PAM (8-11 bps) were used as well, with the same number of bases post-PAM as the sequences without PAM but with an additional 3 (TTN PAM) or 4 (TTTN PAM) nts prior to the 4 to 7 PAM-proximal NT-strand. Searches were performed on both Target Strand and Non-Target Strand, both in 5’→3’ direction.

## Notes

### Competing Interest Statement

Guillermo Montoya declares that he is a co-founder of Twelve Bio

