## Supplementary figures and images for "Cas12a is a dynamic and precise RNA-guided nuclease without off-target activity on λ-DNA"

### Supplementary Figure 1

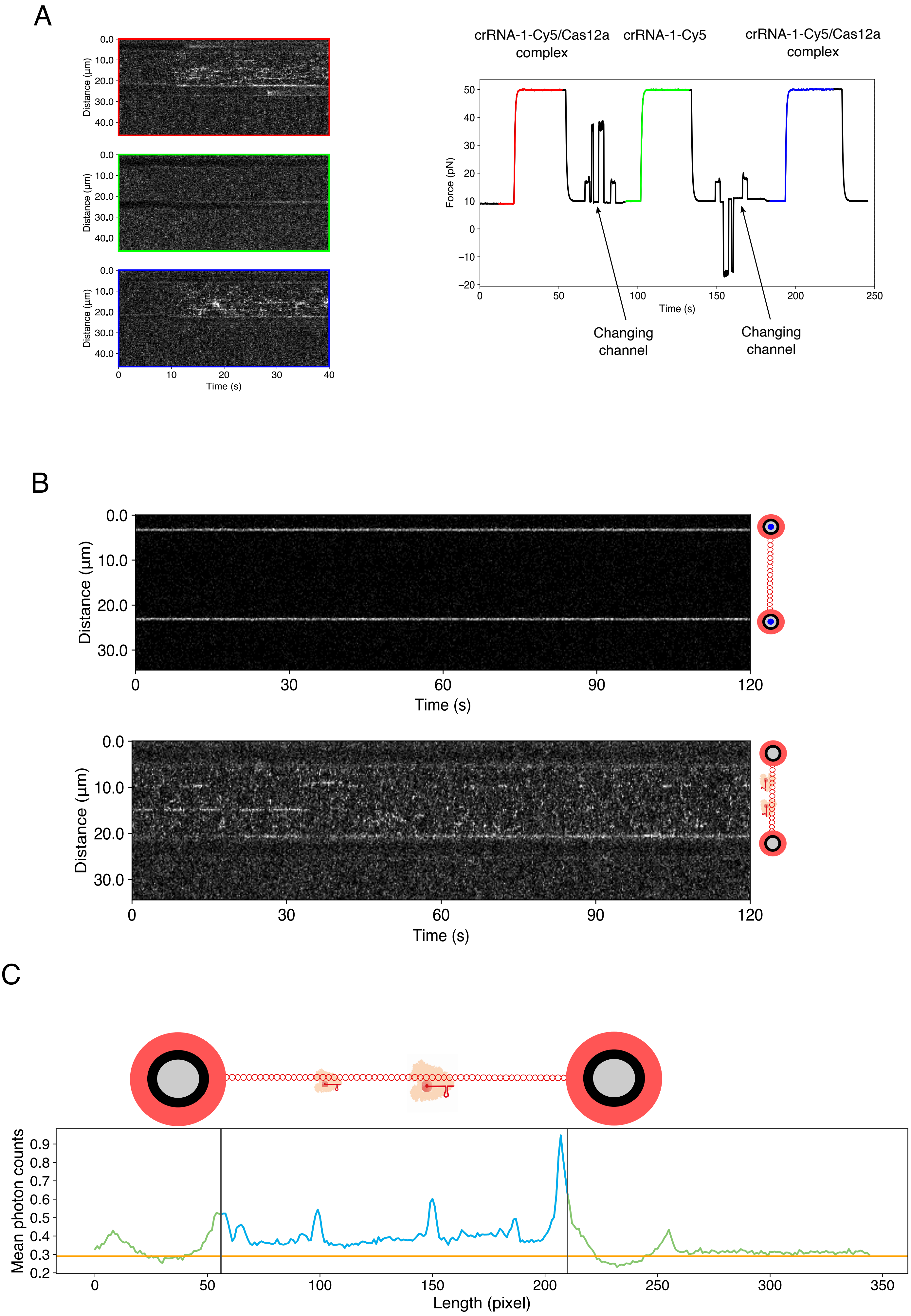

### Supplementary Figure 2

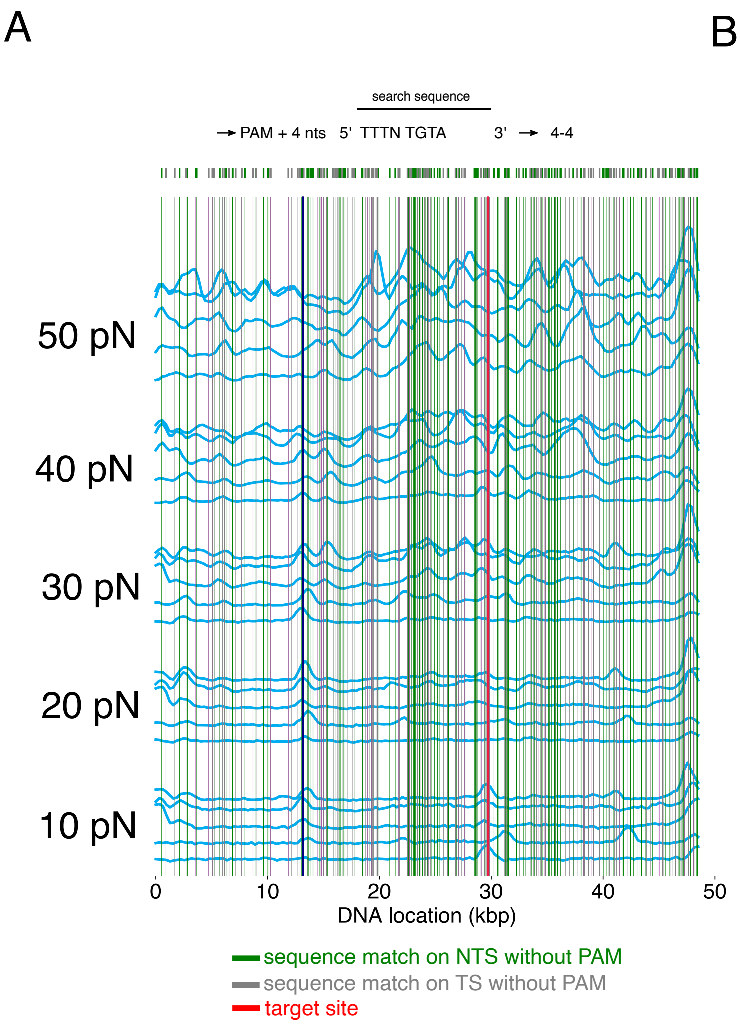

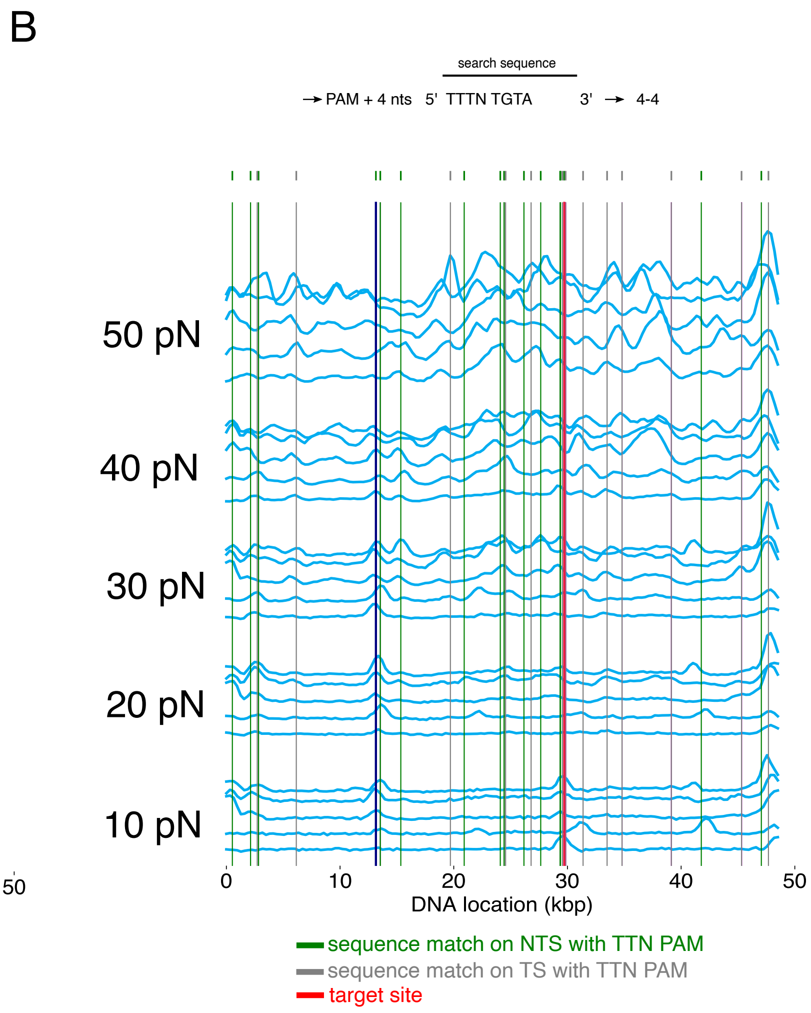

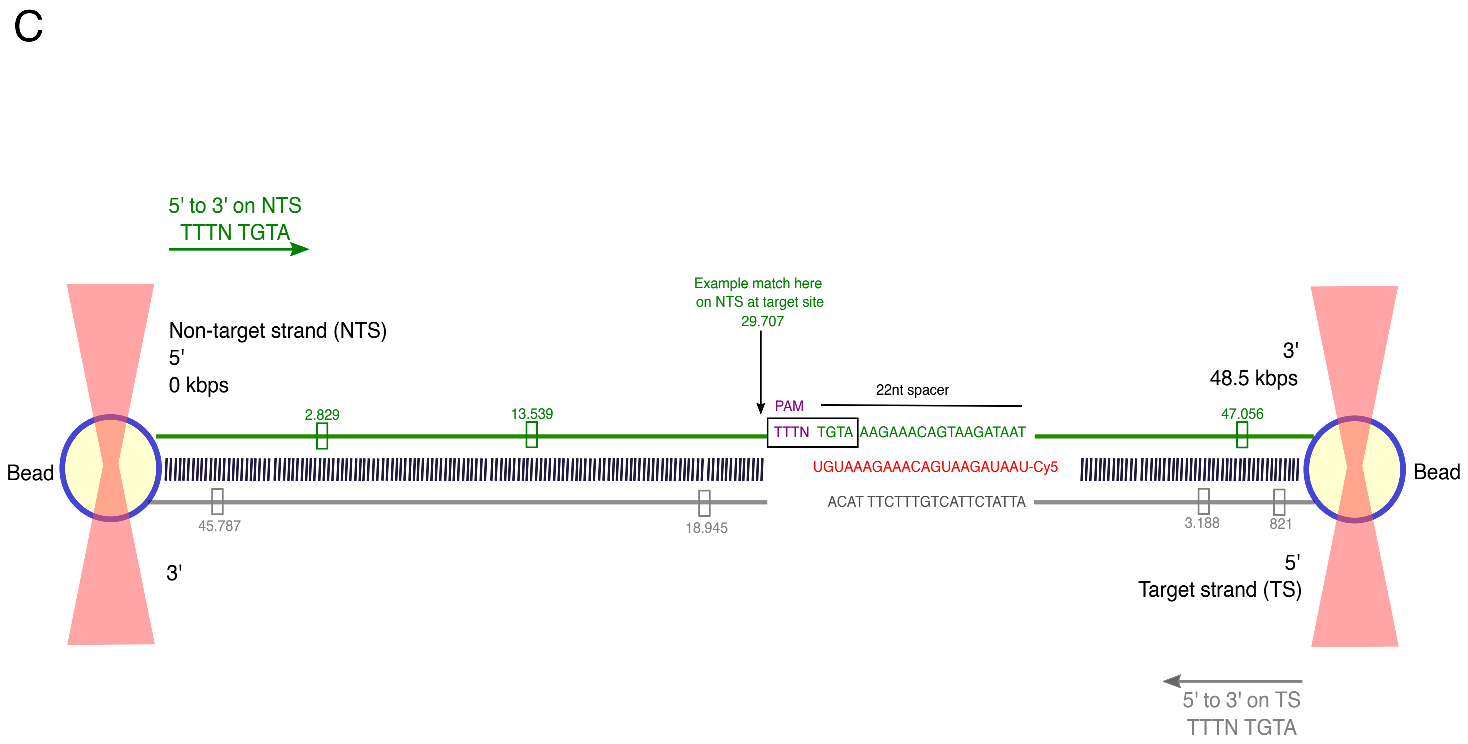

### Supplementary Figure 3

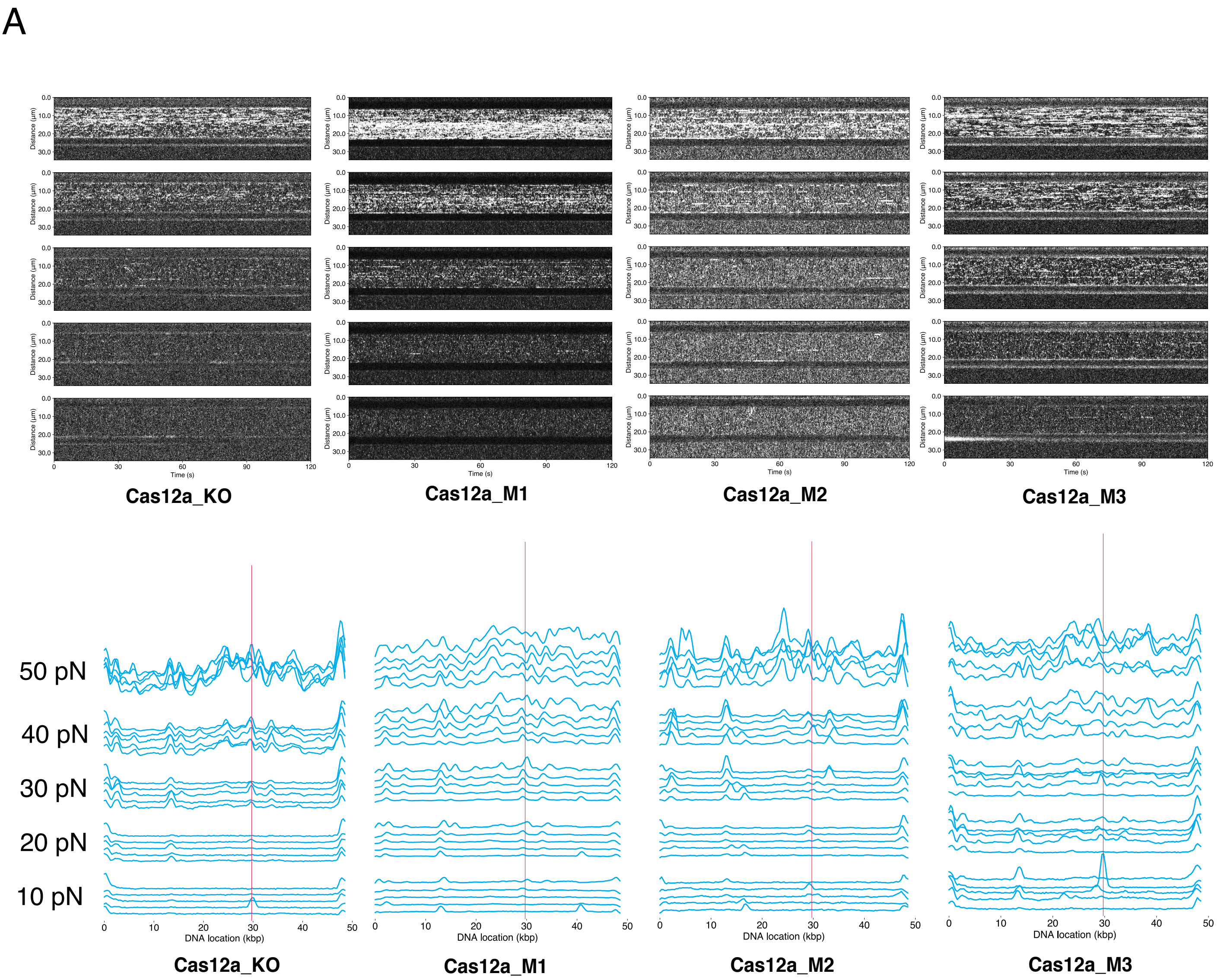

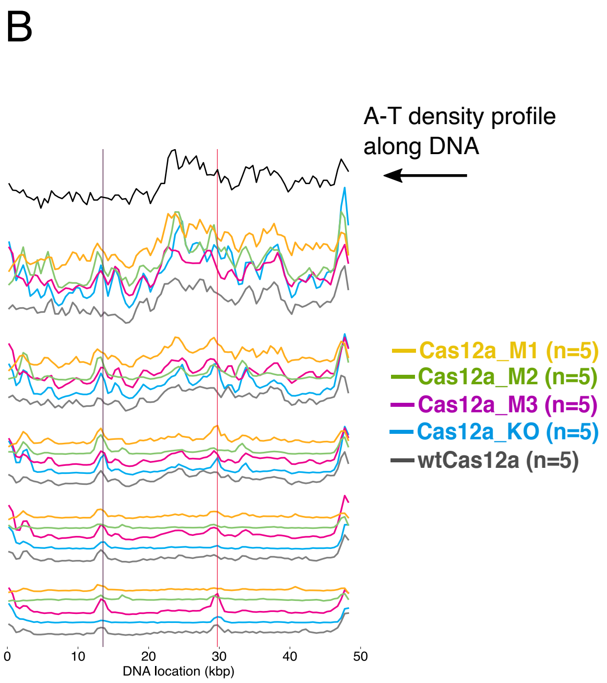

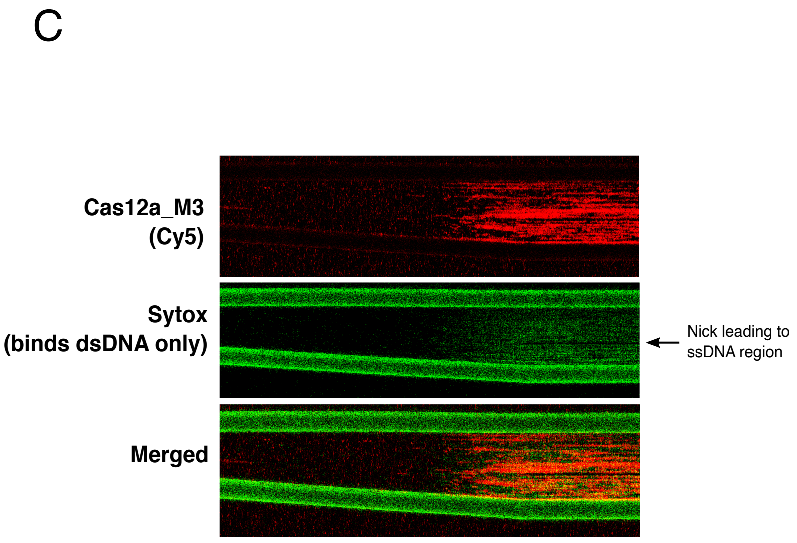

### Supplementary Figure 4

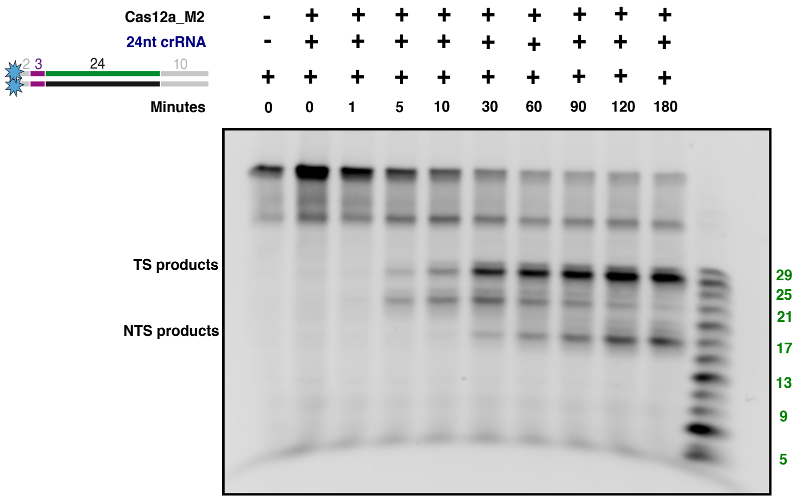

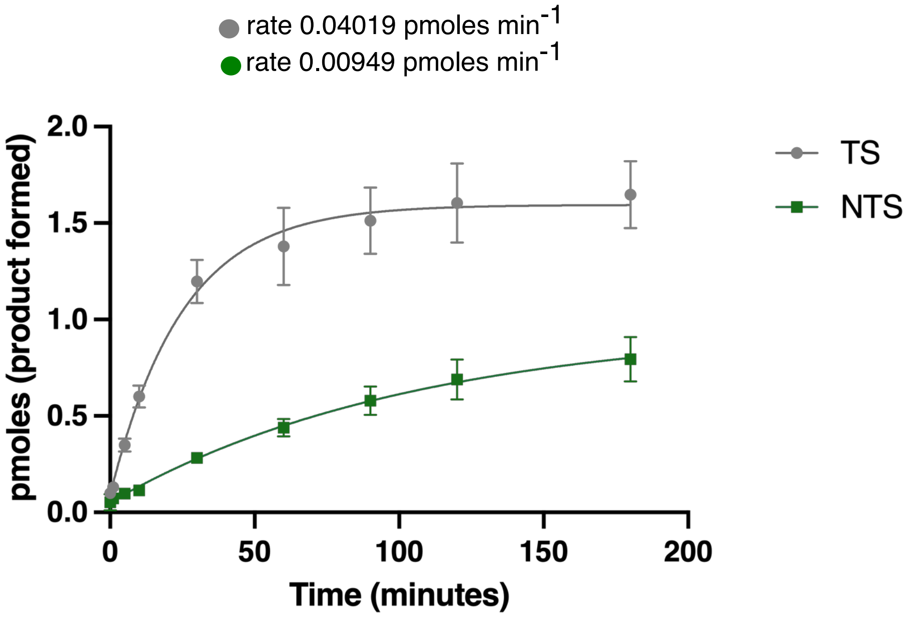
