## Supplementary data table for "Cas12a is a dynamic and precise RNA-guided nuclease without off-target activity on λ-DNA"

| No | Name | Sequence |
| --- | --- | --- |
| 1 | 3´-Cy5-labeled crRNA-1 | rArArUrUrUrCrUrArCrUrGrUrUrGrUrArGrArUrUrGrUrArArArGrArArArCrArGrUrArArGrArUrArArU/3Cy5Sp/ |
| 2 | λ-activator_40nt | CATCGGGTTGAGTATTATCTTACTGTTTCTTTACATAAAC |
| 3 | Unspecific ssDNA | CGATGCATGACCAGTCTTTATCCCCCAGGTCCTCCAAACT/36-FAM/ |
| 4 | crRNA-2 | rArArUrUrUrCrUrArCrUrGrUrUrGrUrArGrArUrGrArGrArArGrUrCrArUrUrUrArArUrArArGrGrCrCrArCrU |
| 5 | TS-FAM labelled | CGATGCATGCAGTGGCCTTATTAAATGACTTCTCTAACG/36-FAM/ |
| 6 | NTS-FAM labelled | /56-FAM/CGTTAGAGAAGTCATTTAATAAGGCCACTGCATGCATCG |
| 7 | Activator TS | CGATGCATGCAGTGGCCTTATTAAATGACTTCTCTAACG |
| 10 | Off target | TCCTGCTCCGGATCGGCGTAACTGTTTCCGTTGACGAAGTTCACCGCATCCAGAAAACGGGCGTAAACCTTACGCCGGACCACCGTTCCGCCGACCAGACTCTGCATATCTTCCGCCATCCCGGTGACCATACCGTACAGGTTAGAAACCGTCAGCGTGGGGCGCGTACTGGTGCCTTTGCCATTCAGTTCAAAACCGCTCCCCTGAATGGGATACGGCTGATACTGTCGCCCCTGCCAGGTGACCGGCTCACCTTTTTCGTTCTGCTCATTACAGAAAAAATAACGTTCTCCACCGACCTCTGTCAGGTCGATTTCCCAGAGCACCACGCTGGCCGACTGCTCCGCACGG |
| 11 | On target | ATGATGTCTGACGCTGGCATTCGCATCAAAGGAGAGTGAGATCGGTTTTGTAAAAGATAACGCTTGTGAAAATGCTGAATTTCGCGTCGTCTTCACAGCGATGCCAGAGTCTGTAGTGTCAGATGATGACCGTACTCAAACATCGGGTTGAGTATTATCTTACTGTTTCTTTACATAAACATTGCTGATACCGTTTAGCTGAAACGACATACATTGCAAGGAGTTTATAAATGAGTATCAATGAGTTAGAGTCTGAGCAAAAAGATTGGGCGTTATCAATGTTGTGCAGATCCGGTGTCTTGTCTCCATGCAGACATCACGAAGGTGTTTATGTAGATGAAGGTATAGAT |
| 12 | λ-DNA | <https://www.ncbi.nlm.nih.gov/nuccore/J02459.1?report=fasta> |
